# m6A methylation dynamically participates in the immune response against *Vibrio anguillarum* in half-smooth tongue sole (*Cynoglossus semilaevis*)

**DOI:** 10.1101/2024.06.23.600224

**Authors:** Suxu Tan, Wenwen Wang, Sen Han, Kunpeng Shi, Shaoqing Zang, Zhendong Wu, Zhenxia Sha

**Affiliations:** Institute of Aquatic Biotechnology, College of Life Sciences, Qingdao University, Qingdao, 266071, China; Laboratory for Marine Fisheries Science and Food Production Processes, Qingdao Marine Science and Technology Center, Qingdao, Shandong 266237, China

**Keywords:** *Cynoglossus semilaevis*, *Vibrio anguillarum*, m6A, epitranscriptome, immune response

## Abstract

N6-methyladenosine (m6A) is the most prevalent RNA modification and a multifaceted regulator capable of affecting various aspects of mRNA metabolism, thereby playing important roles in numerous physiological processes. However, it is still unknown whether, when, and to what extent m6A modulation are implicated in the immune response of an economically important aquaculture fish, half-smooth tongue sole (*Cynoglossus semilaevis*). Herein, we systematically profiled and characterized the m6A epitranscriptome and transcriptome in *C. semilaevis* after the infection of *Vibrio anguillarum*, a serious threat to the aquaculture industry. We demonstrated that m6A could be modulated as early as 4-hour post infection (hpi), and the overall intensity of m6A methylation was enhanced following infection. Both conservative and novel motifs were uncovered from the m6A modification sites. Furthermore, differentially m6A methylated genes (DMGs) and differentially expressed genes (DEGs) were identified, and functional enrichment revealed multiple immune-related pathways, especially the FoxO signaling pathway which showed significance in every comparison. Joint analysis highlighted the remarkedly dynamic role of m6A on gene expression, i.e. early on, m6A mainly prioritized the down-regulation of specific genes, and later, it switched gears to promote expression of another set of genes. Moreover, key immune-related genes, including *pdp1*, *rgs5l*, and *plk2b*, were identified. To our limited knowledge, this is the first study comprehensively characterizing the global m6A atlas in aquaculture fish species. The presented results provide new insights into the dynamics of m6A modifications in the transcriptome of the half-smooth tongue sole following bacterial infection. Further studies are warranted to elucidate the functional significance of these changes and to understand how they affect specific biological processes.

## 1. Introduction

N6-methyladenosine (m6A) methylation is a post-transcriptional modification that occurs on adenosine residues in RNA. First discovered in 1955 (Dunn and Smith, 1955), it has been gradually found to be prevalent from virus, bacteria, yeast to mammals. The identification of methyltransferase METTL3 (Bokar et al., 1997) as well as the discovery of FTO and ALKBH5 as demethylases (Jia et al., 2011; Zheng et al., 2013) are important milestones, showing that m6a is a reversible and dynamic RNA modification. Following these, more regulatory enzymes have been found and catalogued into “writers” which catalyze the addition of the methyl group, “erasers” which remove the modification, and “readers” that recognize and interpret this epigenetic mark. m6A modification has been revealed to impact multiple molecular functions, including RNA transcription, stability, splicing, polyadenylation, localization, and translation (He and He, 2021; Zaccara et al., 2019). As such, a variety of biological processes are regulated by m6A modification, including development, differentiation, diseases, and immune responses.

Studies have demonstrated the involvement of m6A methylation in mammalian immunity (Gan et al., 2023; Liu et al., 2021; Shulman and Stern-Ginossar, 2020), including the regulation of immune cell function and immune responses. m6A modification is widely involved in innate immunity. For instance, after the *Salmonella typhimurium* infection in human macrophages, the m6A reading protein YTHDF2 reduces the mRNA stability of the demethylase gene *KDM6B*, thereby promoting the histone H3K27me3 methylation to reduce the transcription of various pro-inflammatory cytokines to maintain immune homeostasis (Wu et al., 2020). Studies in mice showed that m6A modification mediated by METTL3 enhanced the mRNA translation of genes such as CD40 and CD80 in dendritic cells, which subsequently activated TLR4/NF-κB signaling pathway to induce the production of cytokines (Wang, H. et al., 2019). Moreover, after LPS stimulation or *Escherichia coli* infection in porcine intestinal epithelial cells, YTHDF1 affected the translation of TRAF6 (tumor necrosis factor receptor-associated factor 6) and then activated NF-κB and MAPK signaling to regulate the release of intestinal epithelial cytokines (Zong et al., 2021). Moreover, RNA m6A modification is involved in the adaptive immune response, playing important roles in maintaining the homeostasis and functions of T cells and B cells (Li et al., 2017; Tong et al., 2018; Zheng et al., 2020).

In contrast, the involvement of m6A methylation in fish immunity has been limited. Xu’s group (Geng et al., 2023a; Geng et al., 2022; Geng et al., 2023b) demonstrated that YTHDF2 could degrade mRNA of STING1 (stimulator of IFN genes) and NOD1 (nucleotide-binding oligomerization domain containing 1) to inhibit the innate immune response, while YTHDF1 promoted the translation of myd88 mRNA and enhance the immune response in croaker. Yao et al. observed that METTL3 suppressed viral infection by enhancing type I interferon responses in sea perch (*Lateolabrax japonicus*) (Yao et al., 2023). The focus of these studies primarily revolved around well-established regulators of m6A methylation. In addition, there is a lack of comprehensive profiling and characterization of m6A atlas as well as its integration with transcriptome-wide gene expression in fish immunity. Half-smooth tongue sole (*Cynoglossus semilaevis*) has been a great model for genomic research and an important aquaculture species (Chen et al., 2014). In its aquaculture industry, *Vibrio anguillarum* has long been a common bacterial pathogen that causes high morbidity and mortality, leading to significant economic losses. Furthermore, as a representative of Gram-negative bacterium, it has been widely used to study the fish immunity (Frans et al., 2011). The present study aims to investigate the profile and characterization of m6A methylation during the immune response and its potential effects on the transcriptome of *C. semilaevis* against *V. anguillarum*.

In this study, we found that m6A modification was ubiquitous in half-smooth tongue sole, and identified conserved/novel sequence motifs. m6A profiles changed significantly and dynamically after the bacterial infection, even in the very early stage following infection. Through the integrated analysis of epitranscriptome and transcriptome, we revealed the putative regulatory relationships between m6A and gene expression. Furthermore, key genes and pathways were uncovered, laying foundations for future functional analysis. This study provides valuable insights into the RNA epigenetic regulation of immunity in half-smooth tongue sole, which may contribute to the development of novel strategies for disease prevention and treatment in aquaculture, an increasingly important sector of the global food economy.

## 2. Materials and methods

### 2.1. Experimental fish and sample collection

Half-smooth tongue soles (weight 23 ± 3 g, length 16 ± 2 cm) were purchased from Dachuan Aquatic Co. (Rizhao, China). All fish were acclimated in a 50 L aerated seawater (temperature 22°C, salinity 26‰, dissolved oxygen 6.5 mg/L) for one week. The *V. anguillarum* used in the experiment was confirmed by 16S rDNA sequencing (Tsingke, China). The bacteria were incubated to mid-logarithmic stage at 28°C in 2216E medium (Hopebio, China), and the concentration was determined by the colony-forming unit (CFU) method.

The experimental fish were randomly divided into the control group and treatment group (30 fish per group with three replicates). The infection experiment was performed by intraperitoneal injection. The treatment group were injected with 50 µL *V. anguillarum* at the concentration of 1.3 × 10^8^ CFU/mL. The concentration used was determined by pre-experiment. The control group was injected with the same volume of phosphate buffered saline (PBS) (Solarbio, China). Liver tissues were collected at 0 h, 4 h and 24 h post *V. anguillarum* infection (hpi), and the corresponding groups were designated to Co, V4, and V24, respectively. All samples were snap frozen in liquid nitrogen and then stored at -80°C for RNA extraction and the following experiments.

All the animal experiments were performed in accordance with the guidelines approved by the Ethics Committee of Qingdao University (No. QDU-AEC-2022386).

### 2.2. RNA extraction, cDNA library construction and m6A-seq

Total RNA was isolated using Trizol reagent (Invitrogen, USA) following the manufacturer’s procedure. The RNA amount and purity of each sample was controlled by NanoDrop 1000 (Thermo Scientific, USA) and 1% agarose gel electrophoresis (Tsingke, China), and the RNA integrity was assessed by Bioanalyzer 2100 (Agilent, USA).

mRNA was purified from total RNA using Dynabeads Oligo (dT) magnetic separation technique and fragmented into small pieces (150-200 nt). Every sample for m6A-seq (Dominissini et al., 2013) was catalogued into two libraries. To enrich m6A-modified RNA, the cleaved RNA fragments were incubated with m6A-specific antibody in immunoprecipitation (IP) buffer (50 mM Tris-HCl, 750 mM NaCl and 0.5% Igepal CA-630) for 2 h at 4°C. The IP RNA was reverse transcribed to cDNA by SuperScrip II Reverse Transcriptase (Invitrogen, USA), which was then used to construct the IP library following the protocol (NEB, USA). The input library was constructed without the antibody treatment, which could be used for the control of the IP library and for reflecting the gene expression level. A total of 18 Libraries (3 time points × 3 replicates × 2) were sequenced using Illumina NovaSeq 6000 in PE150 mode (LC-Bio Technology, China).

### 2.3. Analysis of m6A methylation and gene expression

The read quality assessment, adapter trimming, and low-quality read filtering were performed using the all-in-one FASTQ preprocessor, fastp (Chen et al., 2018). The high-quality reads were then aligned to the reference genome of half-smooth tongue sole (Chen et al., 2014) using HISAT2 (Kim et al., 2019).

Calling and differential analysis of m6A read peak were employed using the R package exomePeak (Meng et al., 2014). Their distribution on the functional element (5’UTR, start_codon, CDS, stop_codon, and 3’UTR) were catalogued using ANNOVAR (Wang et al., 2010). HOMER was used to identify the motif after peak calling (Heinz et al., 2010). Visualization of m6A methylation was achieved via R package BioSeqUtils (Zhang, 2023).

Expression level of genes was obtained using StringTie (Pertea et al., 2016). Series Test of Cluster (STC) analysis was performed to classify genes and show the gene expression trend after the bacterial infection. Principle component analysis (PCA) was performed to show the overall variation in gene expression among samples by reducing the dimensionality. Analysis of differential gene expression was performed using DESeq2 (Love et al., 2014). The significance criteria for all differential analyses were set as |log2FC| ≥1 & Q (*P* adjust) value < 0.05.

### 2.4. Functional enrichment analysis

To assign the biological significance, differential genes were then subjected to the enrichment analyses based on terms in GO (Gene Ontology) (2021) and KEGG (Kyoto Encyclopedia of Genes and Genomes) (Kanehisa et al., 2021). The hypergeometric test was employed using the clusterProfiler R package (Yu et al., 2012). The Q value less than 0.05 was used as the threshold for significance.

### 2.5. Joint analysis of m6A methylation and gene expression

Proportional Venn diagram (Lyu et al., 2023) was used to exhibit the common genes that were both differentially m6A-modified and differentially expressed. Four-quadrant diagram was used to show their distribution and relationship of these two datasets, noting that each point represents a peak, not a gene. In addition, common genes underwent protein-protein interaction (PPI) analysis to reveal their interactions and further screen candidate genes (Szklarczyk et al., 2023).

### 2.6. qRT-PCR validation

Quantitative reverse transcription polymerase chain reaction (qRT-PCR) was performed on nine randomly selected DEGs, i.e., three for each comparison. Details of primers are shown in Table 1. cDNA synthesis was conducted by Hiscript III RT Super Mix for qPCR (+gDNA Wiper) reverse transcription kit (Vazyme, China). The expression of gene was evaluated by qRT-PCR on a Borges 9600Plus fluorescent quantitative PCR instrument following the protocol of ChamQ SYBR Color qPCR Master Mix kit (Vazyme, China). The reaction system was configured on ice according to the instructions, and the reaction procedure was as follows: 95°C for 30 s; 95°C for 10 s, 60°C for 30 s, 40 cycles; 95°C for 15 s; 60°C for 1 min and 95°C for 15 s. Relative gene expression was analyzed by 2^-ΔΔCt^. The 18S rRNA gene was used as the internal reference gene, as previously described (Liu et al., 2023; Wang, W. et al., 2023).

**Table 1.**
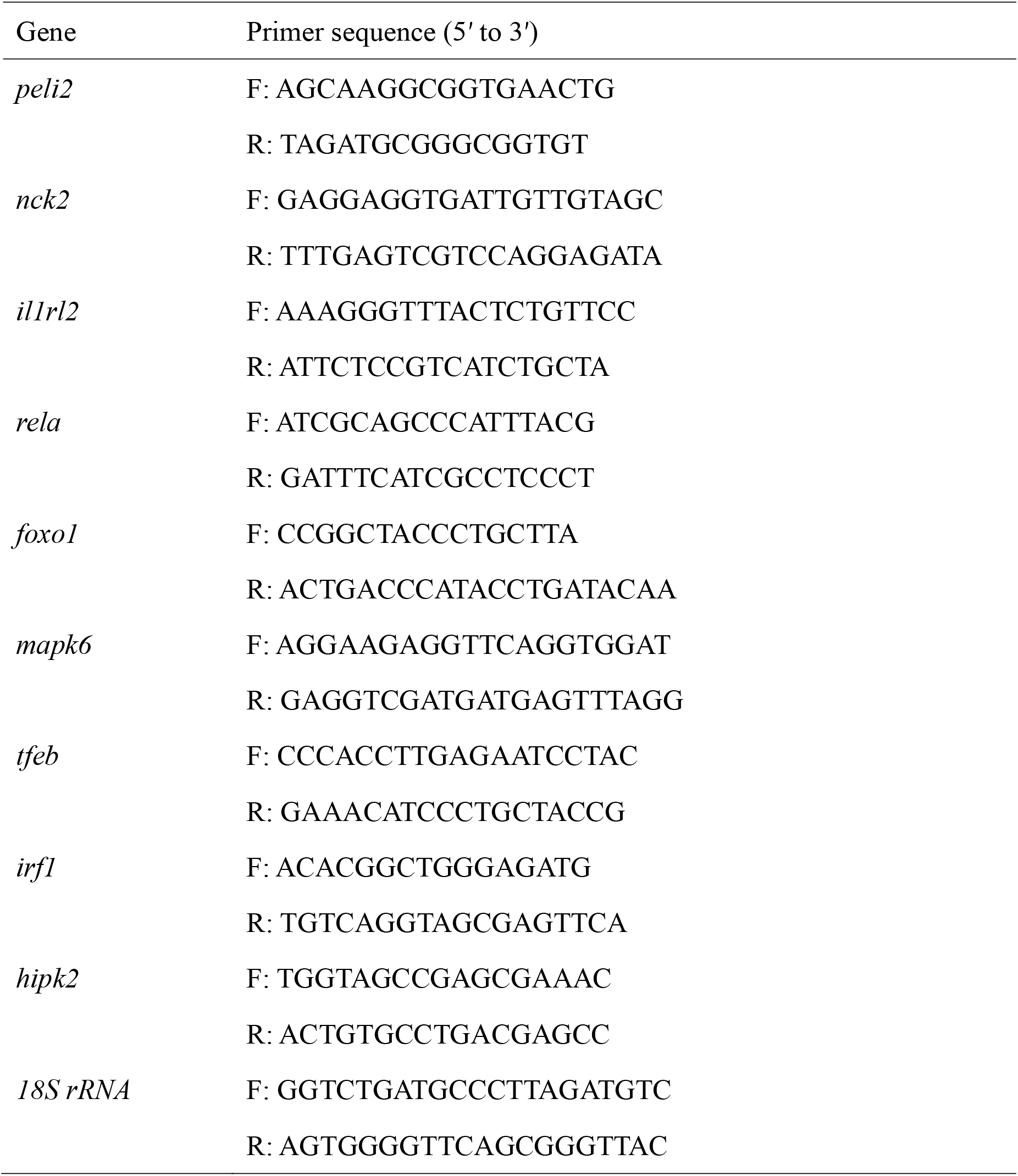
Sequences of the primers used in this study.

## 3. Results

### 3.1. Characterization of m6A methylation

Sequencing data generated in this study are available at the NCBI Sequence Read Archive (http://www.ncbi.nlm.nih.gov/sra) under the accession number PRJNA1109439. Detailed statistics of sequencing and mapping are provided in Table 2. A total of 23,945 m6A peaks were identified in 12,213 genes in the control group. Similarly, a set of 22,685 and 27,186 peaks were found in 12,068 and 13,681 genes in the V4 and V24 group, respectively. On average, one gene had two m6A read peaks. As shown in Figure 1A-C, about half of these m6A modified genes harbor one peak, and most genes have 1-3 peaks. It is noted that each group was calculated with three samples combined. We further characterized the peak distribution on RNA functional elements. The samples at three groups were similar (Fig. 1D-F), with the coding sequence (CDS) containing the highest number of peaks, followed by stop codon, 3’ UTR, and start codon, while 5’ UTR had the lowest number of peaks. In terms of peak density, increased m6A peak density was observed in stop codon and near start codon for all three groups (Fig. 1G). Moreover, the overall level of m6A was detected to explore the impact of bacterial infection. The CDF (cumulative distribution function) was plotted (Fig. 1H), where the X-axis represents the peak enrichment (log2FC) in the peak calling results, and the Y-axis indicates the cumulative percentage of m6A modification. Under the same m6A cumulative percent, the larger the log2FC, the higher the methylation level of the sample. Both the CDF plot and violin plot (Fig. 1H-I) illustrated that the overall m6A level increased after the *V. anguillarum* infection in half-smooth tongue sole (Fig. 1H-I). No evident difference of m6A level was found between samples at 4 hpi and 24 hpi.

**Fig. 1.**
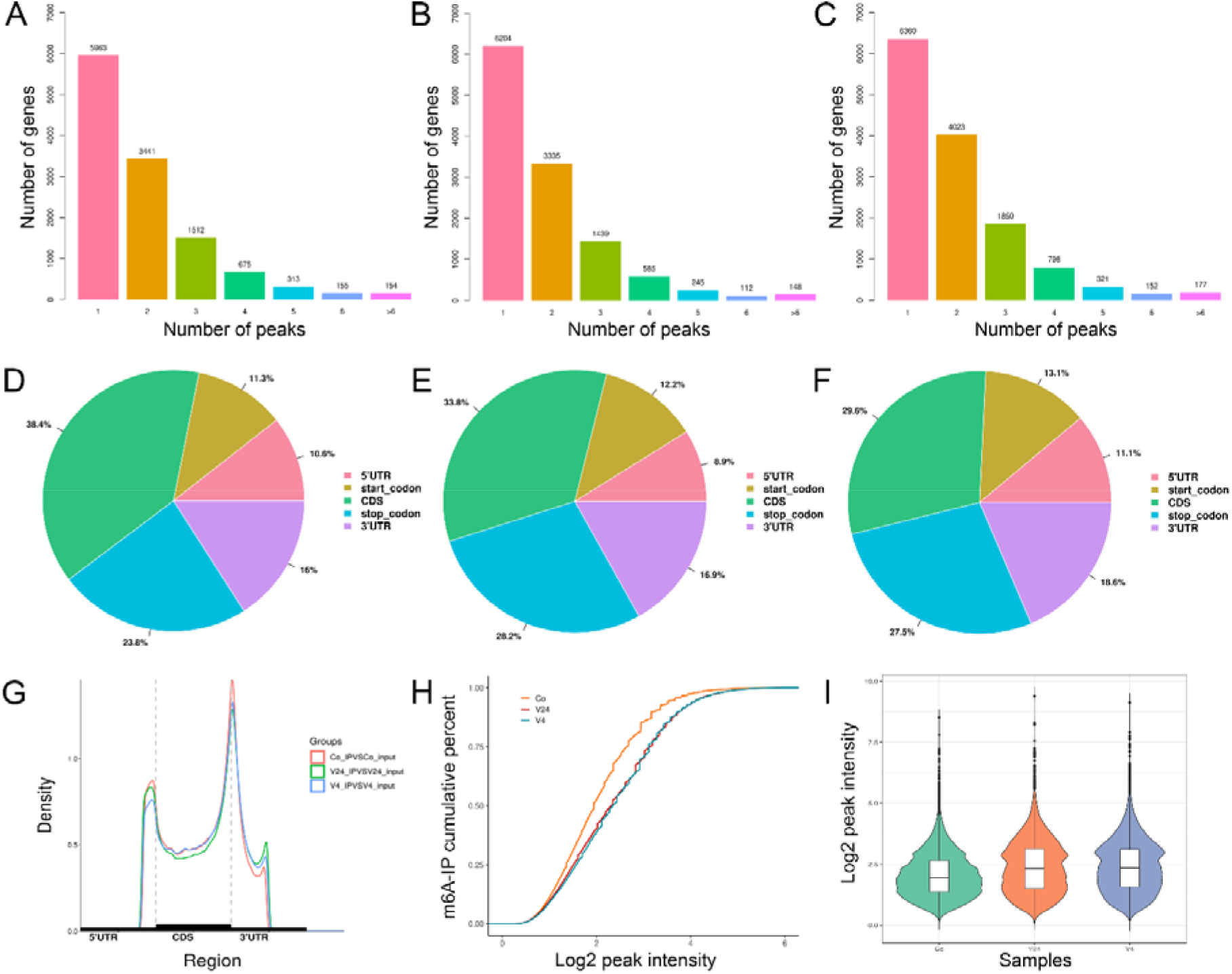
Profiles of m6A methylation in half-smooth tongue sole after infection with *Vibrio anguillarum*. (A-C) The number of m6A genes with respect to the number of m6A peaks per gene in the group of Co (A), V4 (B), and V24 (C). (D-F) The distribution of peaks in different genomic features in the group of Co (D), V4 (E), and V24 (F). (G) The density of m6A peaks in different genomic features in three groups. (H-I) The overall m6A peak density across the epitranscriptome in three groups illustrated by CDF plot (H) and violin plot (I). Co, control; V4, 4 h after the infection; V24, 24 h after the infection.

**Table 2.**
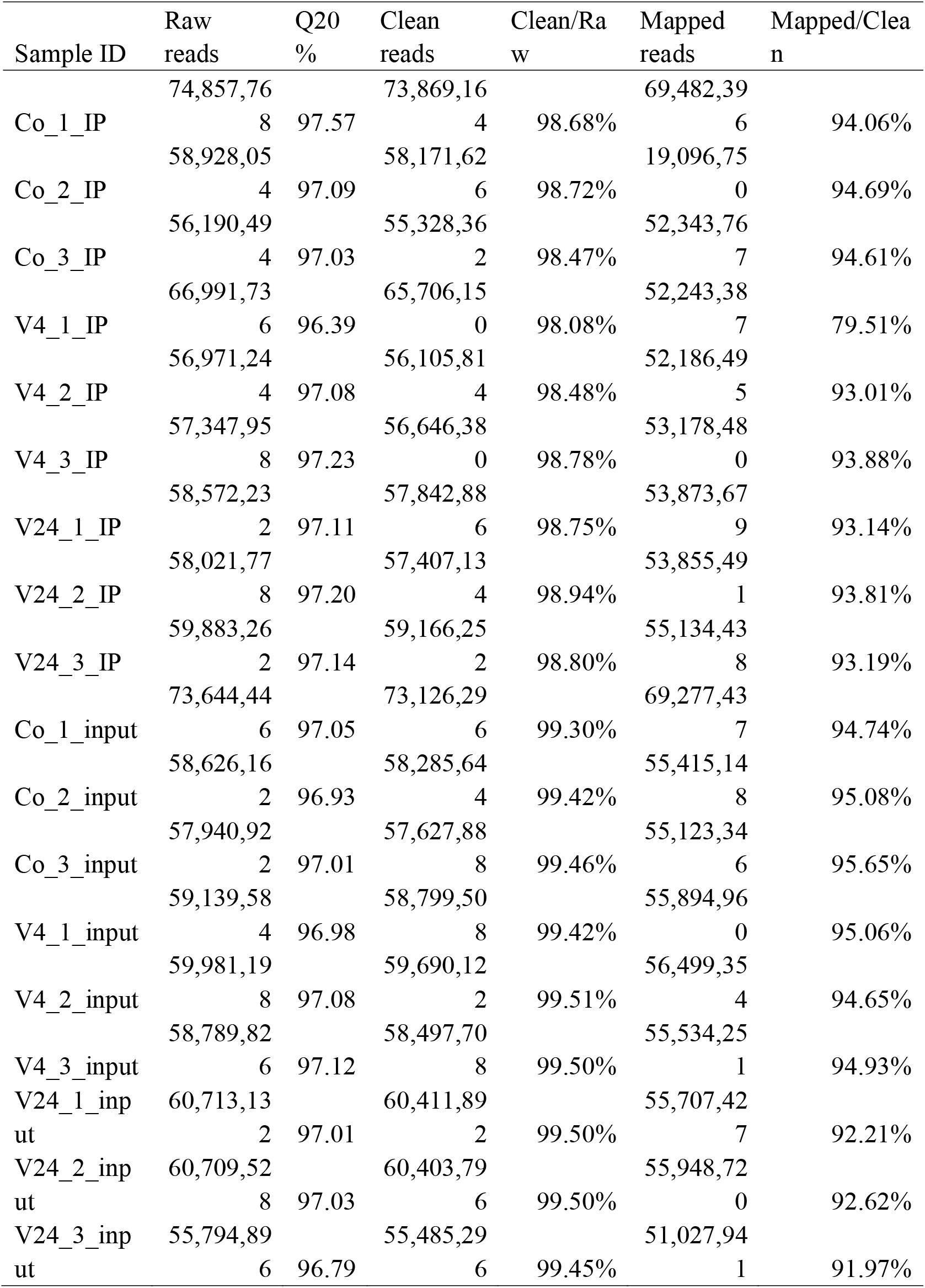
Detailed statistics of sequencing and mapping. Co, control; V4, 4 h after the infection; V24, 24 h after the infection; IP, m6A immunoprecipitation before sequencing; input, RNA-seq without m6A sequencing.

To reveal the characteristics of sequences with m6A modification, i.e. sequences that are specifically combined with methyltransferase, we performed motif analysis. The top three enriched motifs were shown for samples at each time point, including 0 h (Fig. 2A-C), 4 h (Fig. 2D-F) and 24 h post infection (Fig. 2G-I). Not surprisingly, we identified commonly occurring DRACH (D = A, G or T; R = A or G; H = A, C or U) consensus sequence in samples at each time point (Fig. 2A, F, G). In addition to the canonical motif, we identified multiple motifs that are significantly enriched in the IP read peak compared with the input read background, including motifs containing GAA (Fig. 2B, 2E, and 2I) and GGAG (Fig. 2C and 2F).

**Fig. 2.**
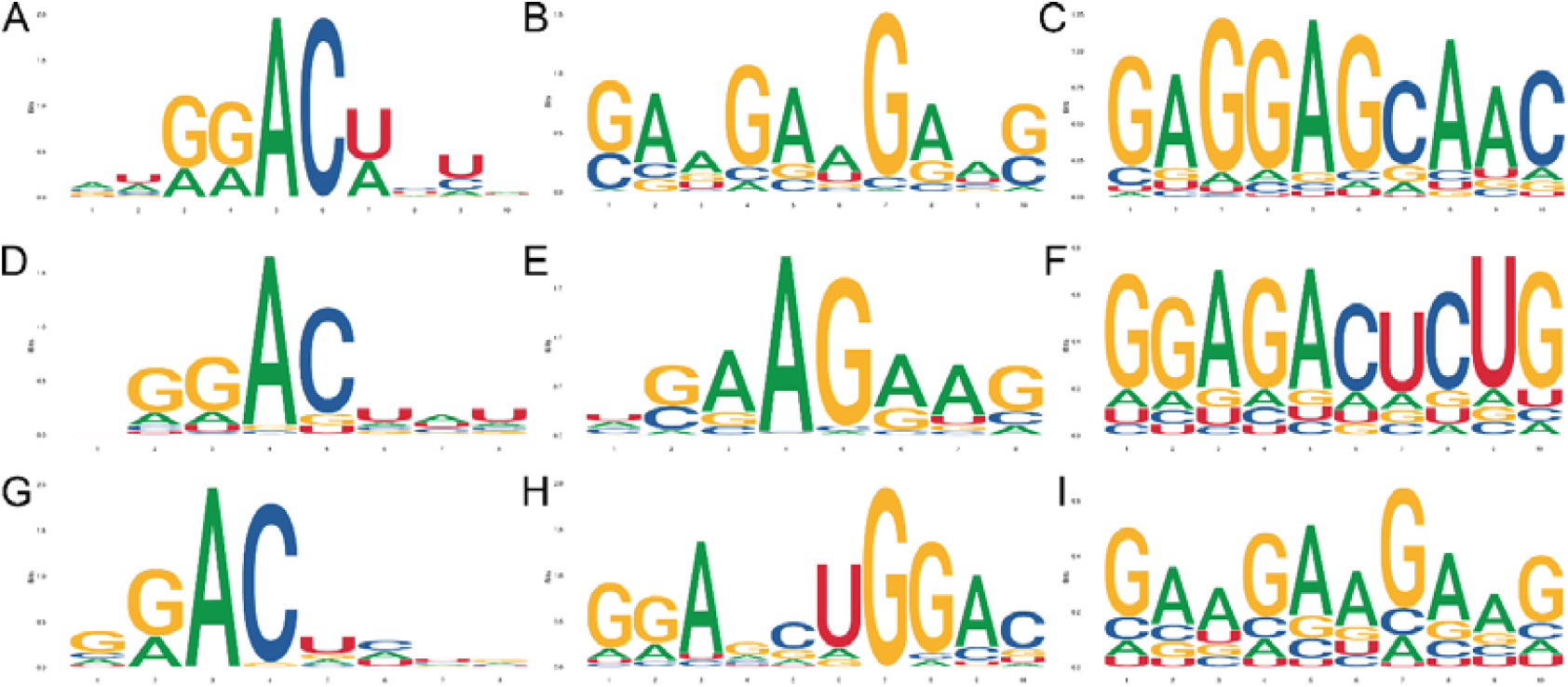
Enriched motifs for m6A methylation regions in the group of Co (A-C), V4 (D-F), and V24 (G-I). Co, control; V4, 4 h after the infection; V24, 24 h after the infection.

### 3.2. Characterization of gene expression

To reveal the gene expression trend and provide new biomarkers related to diseases after bacterial infection, Series Test of Cluster analysis was performed. All expressed genes were grouped into 8 clusters. Cluster 1, 5, and 8 exhibited similar trends, where their expressions were first down-regulated, then up-regulated after the bacterial infection. However, the degree of trend varies. For example, the expression at 24 hpi was higher than that at 0 h for cluster 1, while the expression at 24 hpi was lower than that at 0 h for cluster 8. In contrast, the expression in cluster 2 and 6 were first up-regulated, then down-regulated after the bacterial infection. In cluster 3 and 4, the overall trend was down-regulated, but in different patterns. Whereas the overall expression in cluster 7 was up-regulated. Considering the gene expression trend, candidate marker DEGs of interest in disease prevention/control could be further selected. Genes in each cluster are provided in Table S1.

The principal coordinate analysis (PCoA) was also performed based on expression of all genes, which shows that three treatments were separated clearly and three replicates of each treatment were grouped together, indicating that there are obvious differences in gene expression among different time points after the infection (Fig. 4). At the same time, these results demonstrate the accuracy of this experiment and the rationality of the experimental design.

**Fig. 3.**
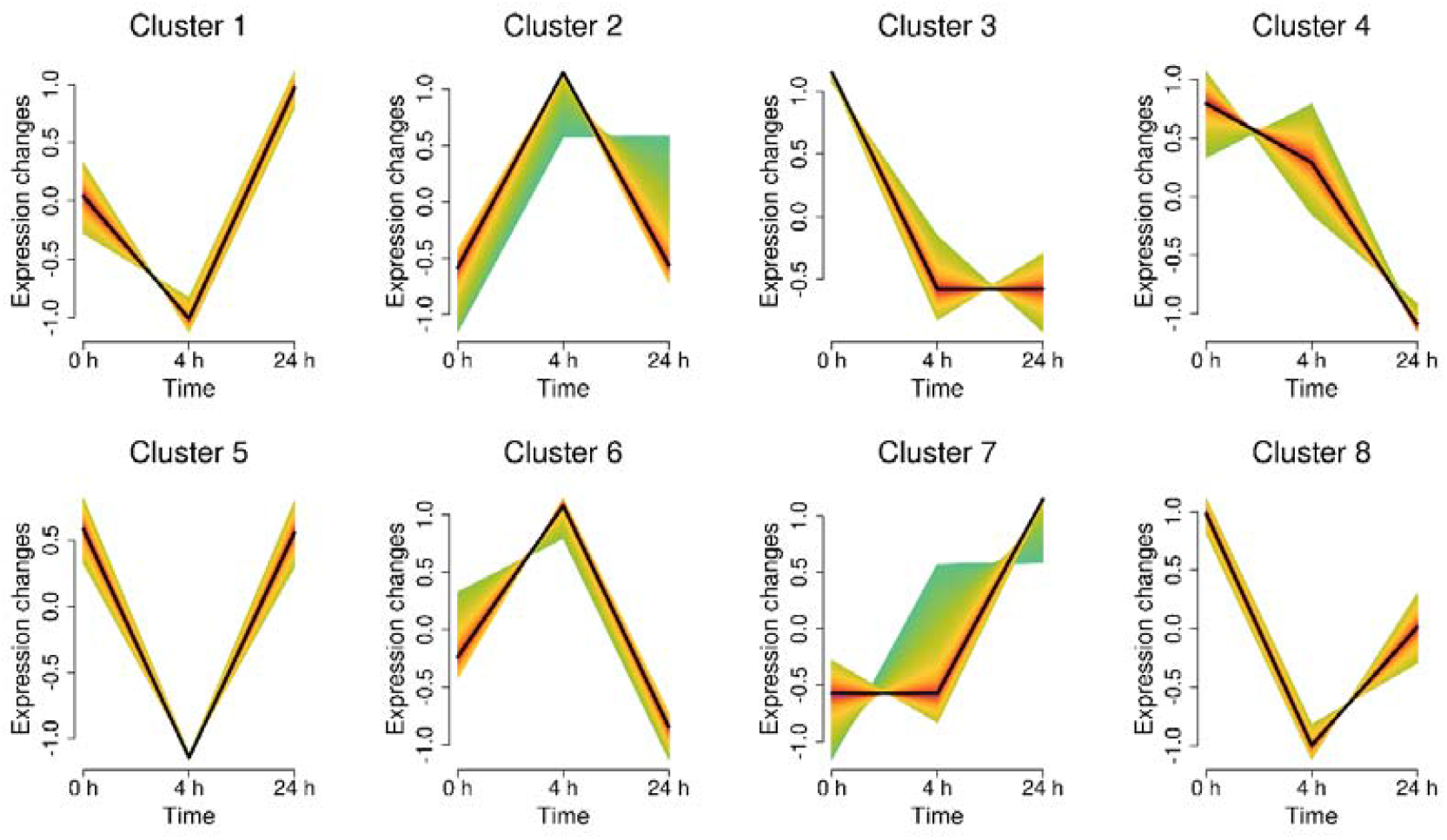
Clustering of genes based on the trend of gene expression in half-smooth tongue sole after infection with *Vibrio anguillarum*.

**Fig. 4.**
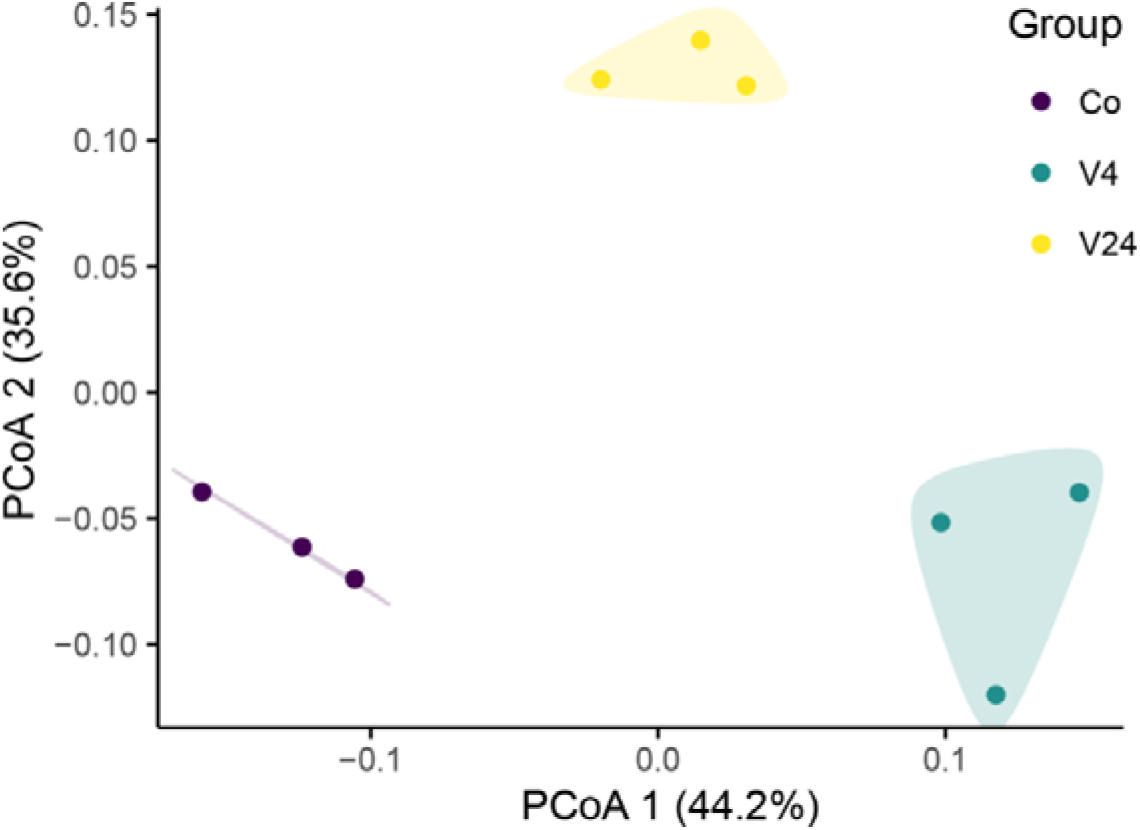
PCoA plot showing the clustering of samples with three replicates in each group. Co, control; V4, 4 h after the infection; V24, 24 h after the infection.

### 3.3. Differentially m6A methylated genes (DMGs) and differentially expressed genes (DEGs)

A total of 302, 970, and 246 differential m6A peaks were identified (Q < 0.05), corresponding to 294, 929, and 240 DMGs in three respective comparisons (V4 vs Co, V24 vs Co, and V24 vs V4). The comparison of V24 vs Co contains the highest differential m6A peaks and genes, suggesting m6A modification accumulates over time following the bacterial infection. The Venn diagram (Fig. 3A) shows the overlapping and specific DMGs among the three comparisons. The three shared genes include *pdp1*, *rgs5l* (*LOC103395811*), and one uncharacterized gene (*LOC103388921*).

In terms of differential expression, a total of 4319, 1691, and 5418 DEGs were identified in the three comparisons (Q < 0.05). Different from DMGs, the comparison of V24 vs V4 contains the most DEGs. The V4 vs Co comparison has more DEGs than the V24 vs Co comparison, and it is noted that these two comparisons also share 451 DEGs which are not shown in the proportional Venn diagram. Most DMGs were specific to the comparison of V24 vs Co, and most DEGs were overlapped between the two infection stages (0-4 hpi and 4-24 hpi). A total of 220 DEGs were shared among the three comparisons (Table S2). No gene was shared in all six comparisons, but 11 genes were shared in five comparisons (Table 3), and most of them are immune-related.

**Table 3.**
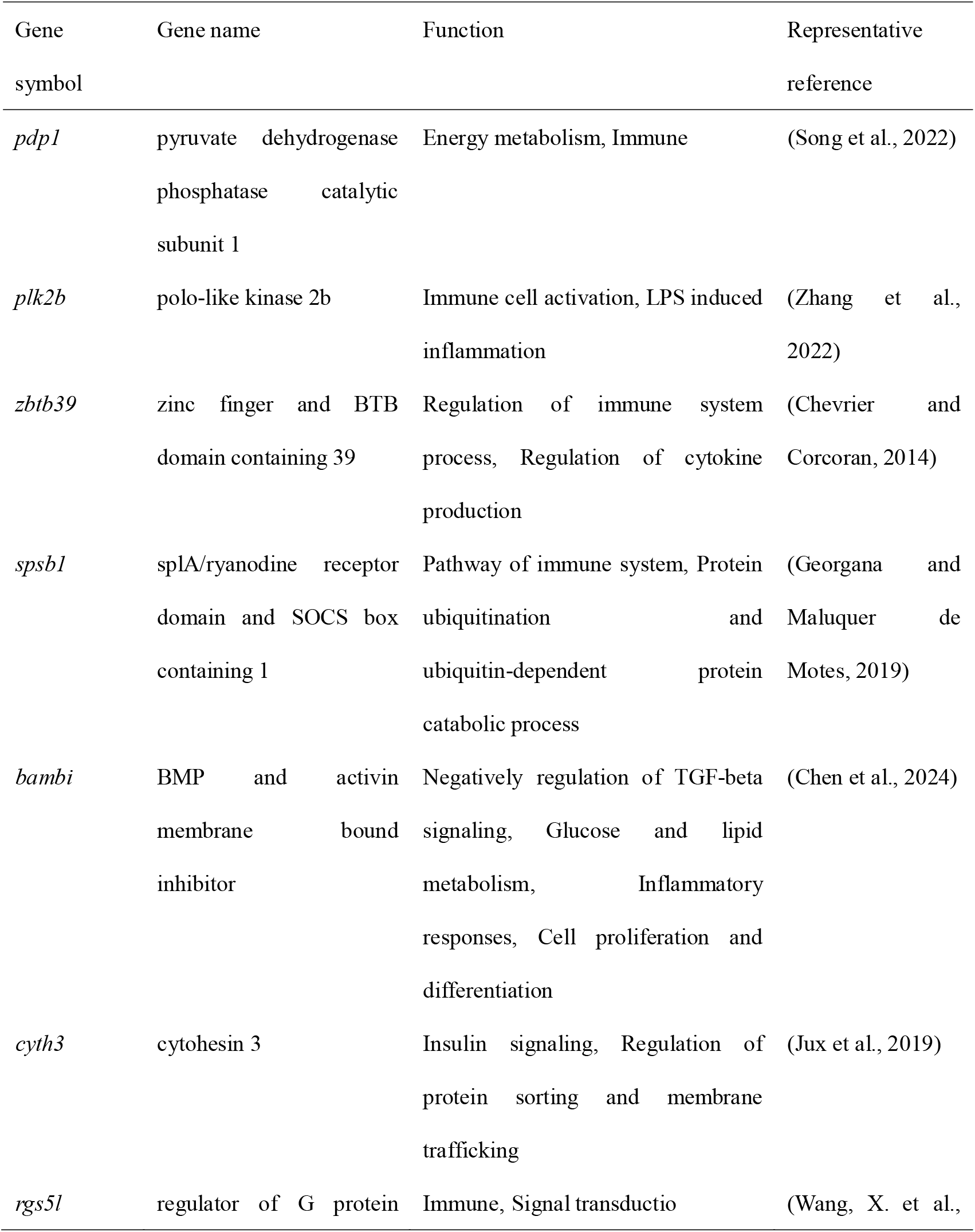
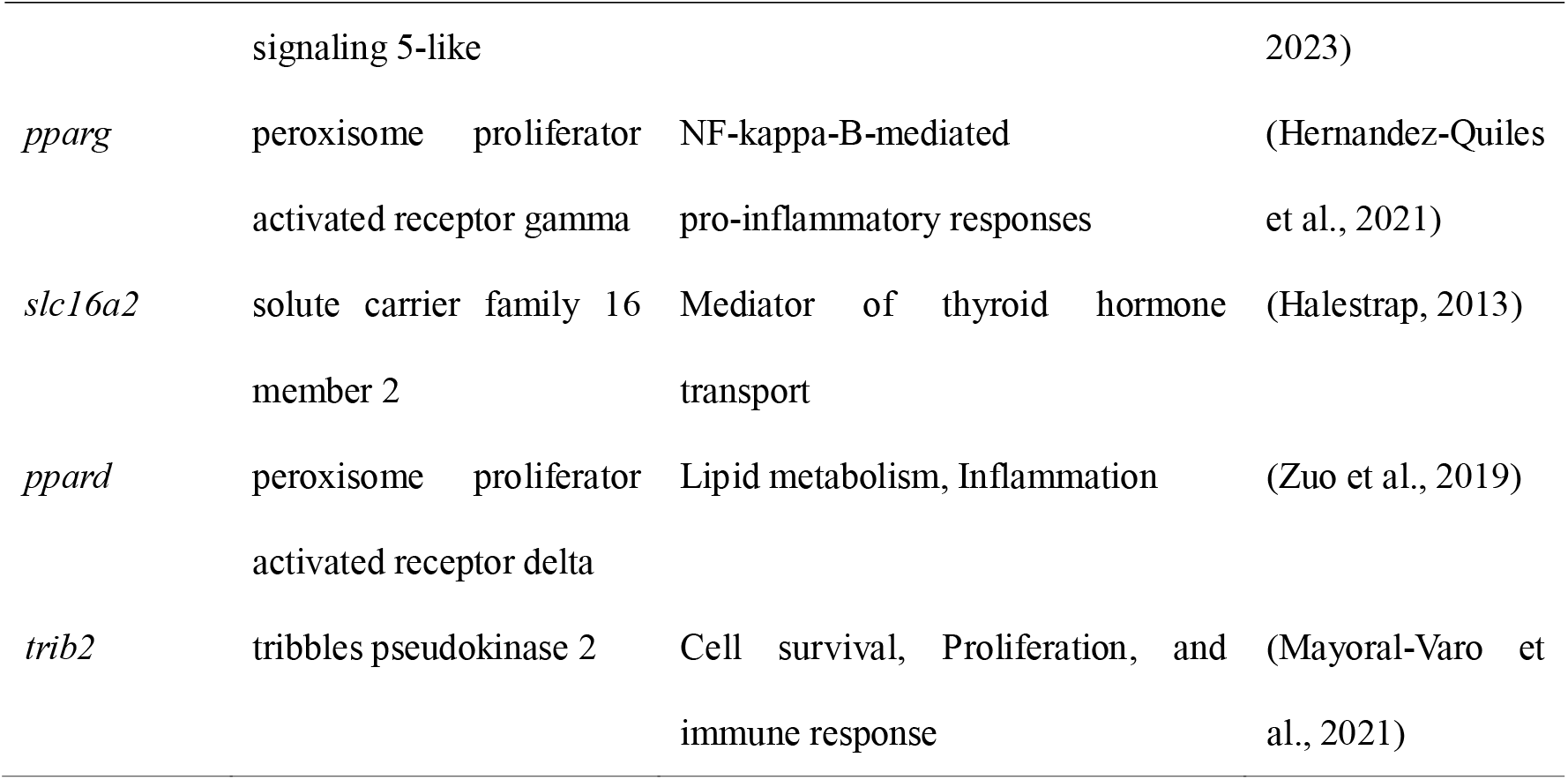
Details of the most common differential genes (DMGs and DEGs) identified in this study. Gene functions were mainly collected form references, UniProt (http://www.uniprot.org), GeneCards (https://www.genecards.org), and Reactome (https://reactome.org/). Except for *cyth3* and *slc16a2*, which are related to energy metabolism, all genes exhibit immune-related functions.

### 3.4. Functional enrichment for DMGs and DEGs

To reveal the functional significance of differential genes, functional enrichment analysis was conducted. We first revealed the enrichment of DMGs. In V4 vs Co comparison, no GO biological process was enriched, but a set of seven pathways were enriched, including MAPK signaling, FoxO signaling, Ras signaling, and glycosaminoglycan biosynthesis, neuroactive ligand-receptor interaction, base excision repair, and purine metabolism pathway, first four of which are immune related (Fig. 6A). In the comparison of V24 vs Co, two GO biological processes were enriched, including regulation of transcription and angiogenesis. A total of 50 signaling pathways were significantly enriched, and multiple pathways are immune-related including but not limited to MAPK signaling, endocytosis, FoxO signaling, mTOR signaling, TGF-beta signaling, PPAR signaling, Glycosaminoglycan biosynthesis, and cytokine-cytokine receptor interaction signaling pathways (Fig. 6C). In V24 vs V4 comparison, we found no significantly enriched GO terms but a total of 11 enriched pathways, including FoxO signaling, endocytosis, PPAR signaling, O-glycan biosynthesis, and amino acid metabolism signaling pathway (Fig. 6E).

**Fig. 5.**
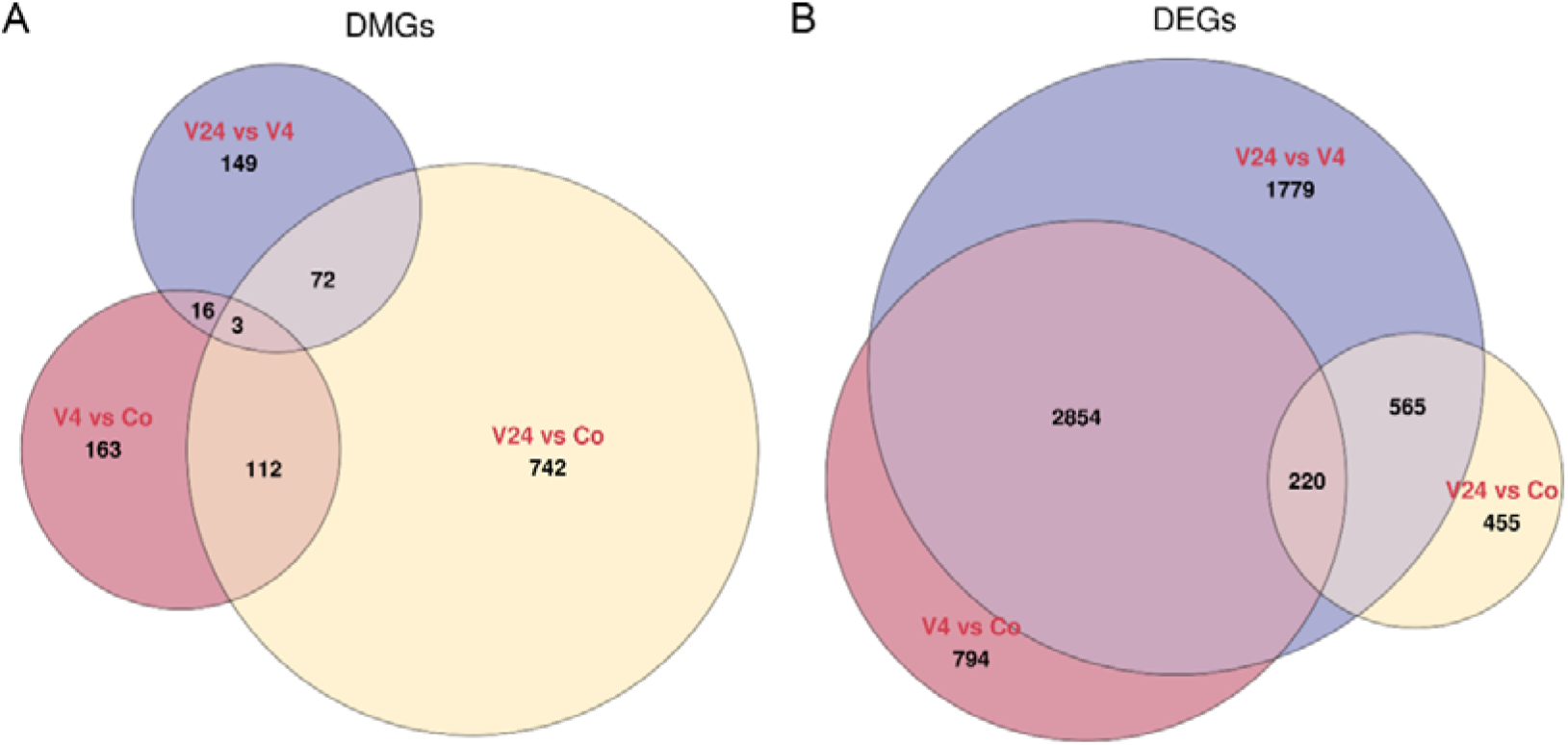
Venn diagram for DMGs (A) and DEGs (B) in half-smooth tongue sole after infection with *Vibrio anguillarum*. DMGs, differentially m6A methylated genes; DEGs, differentially expressed genes; Co, control; V4, 4 h after the infection; V24, 24 h after the infection.

**Fig. 6.**
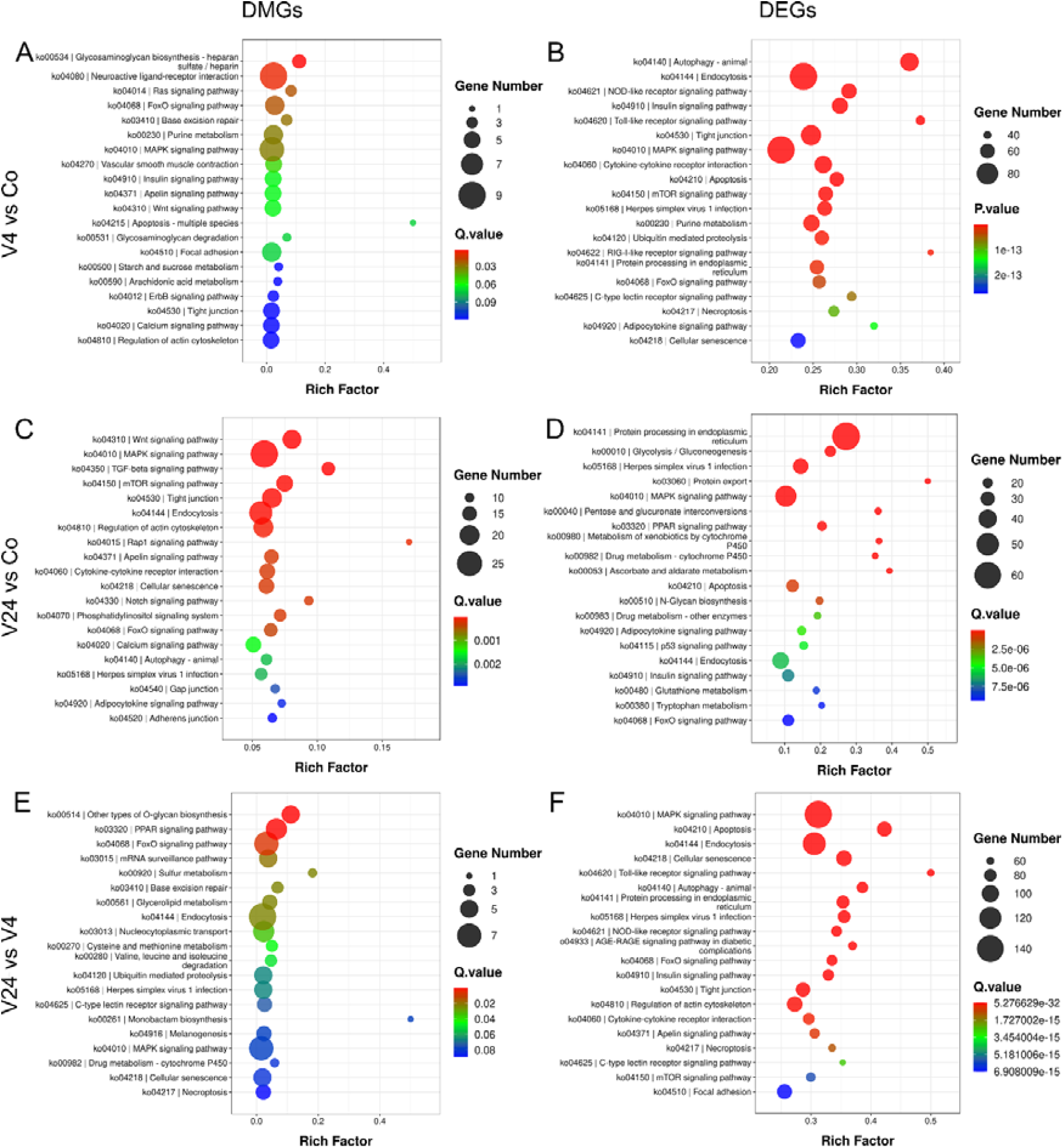
Enriched pathways for DMGs (A, C, E) and DEGs (B, D, F) in half-smooth tongue sole after infection with *Vibrio anguillarum*. DMGs, differentially m6A methylated genes; DEGs, differentially expressed genes; Co, control; V4, 4 h after the infection; V24, 24 h after the infection.

Furthermore, the enrichment of DEGs were studied. In the comparison of V4 vs Co, the enriched GO biological processes include phosphorylation, regulation of transcription, triglyceride homeostasis, and steroid hormone mediated signaling pathway. A total of 142 pathways were significantly enriched (Fig. 6B). Most of the top 20 enriched ones are immune-related, such as autophagy, endocytosis, NOD-like receptor signaling, Toll-like receptor signaling, MAPK signaling, cytokine-cytokine receptor interaction, mTOR signaling, RIG-I-like receptor signaling, FoxO signaling, C-type lectin receptor signaling, and Adipocytokine signaling pathways. Notably, in the same comparison, all seven enriched pathways in DMGs were also enriched in DEGs, highlighting their importance in the early immune response to bacterial infection. In the comparison of V24 vs Co, a set of 23 biological processes were significantly enriched, including response to lipopolysaccharide, response to cytokine, response to cAMP, steroid hormone mediated signaling, response to hormone stimulus, regulation of apoptotic process, oxidation-reduction process, protein folding, and metabolic processes, many of which are closely related to immune response. A total of 95 KEGG pathways were significantly enriched, again, many of which are immune related. For instance, among the top 20 enriched pathways, multiple pathways are involved in immune response, including MAPK signaling, PPAR signaling, metabolism of xenobiotics by cytochrome P450, apoptosis, adipocytokine signaling, p53 signaling, endocytosis, and FoxO signaling pathways (Fig. 6D). It is noted that a total of 30 pathways were shared when comparing the enriched pathways of DEGs and DMGs in the same comparison. We next investigate the enrichment in the comparison of V24 vs V4, a set of 21 biological processes were significantly enriched, including response to lipopolysaccharide, response to cytokine, steroid hormone mediated signaling, protein folding, inflammatory response, negative regulation of I-kappaB kinase/NF-kappaB signaling, cell redox homeostasis, cell migration, and endosomal transport, first four of these immune-associated ones were also enriched in the V24 vs Co comparison. A total of 152 pathways were significantly enriched, and we observed many immune-related pathways; for instance, most of the top 20 enriched ones are immune-related and have been enriched in other comparisons (Fig. 6F).

Intriguingly, FoxO signaling was the only enriched pathway common in all six comparisons (Table S3). The other six enriched pathways were shared in five comparisons, including MAPK signaling pathway, neuroactive ligand-receptor interaction, base excision repair, PPAR signaling pathway, mRNA surveillance pathway, and endocytosis.

### 3.5. Joint analysis of differential m6A modification and differential expression

To intuitively show the relationship and the overall distribution between m6A modulation and gene expression, Venn diagram and four-quadrant diagram were plotted in this study. A set of 131, 122, 96 genes were respectively common differential genes in the comparison of V4 vs Co, V24 vs Co, and V24 vs V4 (Fig. 7 A, C, and E). More than 40 percent of DMGs in the relative early (V4 vs Co) and later stage (V24 vs V4) after the infection contribute to the differential gene expression, while, surprisingly, only two common genes (*chchd2*, *mbd4*) were shared between the early and later stage after the infection, suggesting the dynamic and diverse regulatory effect of m6A which could exert regulation adjustment as the bacterial disease progresses. The common differential genes for each comparison are listed in Table S4. Four-quadrant diagram shows that, m6A methylation is regulated to mainly decrease gene expression during the early stage (V4 vs Co), whereas m6A may be regulated to increase specific gene expression in the relative later stage (V24 vs V4) (Fig. 7 B and F), reinforcing the dynamic regulatory role of m6A. From the overall process (V24 vs Co), m6A methylation mainly plays the role of inhibiting expression (Fig. 7 D).

**Fig. 7.**
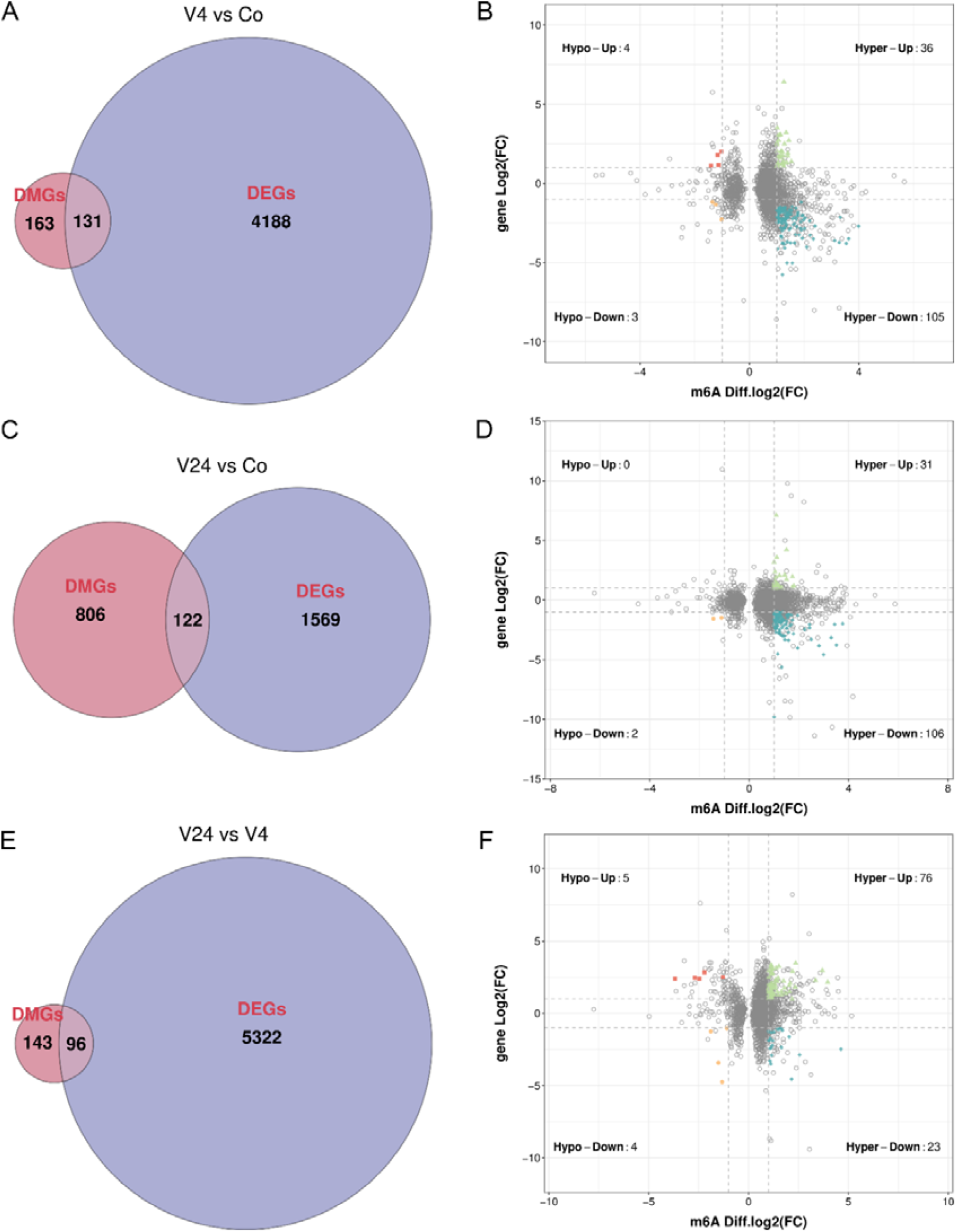
Venn diagram (A, C, E) and four-quadrant diagram (B, D, F) showing the relationship between DMGs and DEGs. In the four-quadrant diagram, each dot represents a m6A peak. The X-axis represents the changes in the methylation modification between groups, and the Y-axis represents the changes in the expression level of the gene where the peak is located between groups. The colored dots in the figure indicate genes with significant differences in both transcription and methylation levels. They are divided into four quadrants: Hypo-up, Hypo-down, Hyper-up, and Hyper-down. The numbers in the quadrants represent the number of peaks in the quadrant. Gray dots represent data with no significant differences. The dotted line represents the log2FC threshold set for the two groups of data.

To reveal the interactions between common genes and finely selecting hub genes, protein-protein interaction analysis was performed for each comparison. As shown in Fig. 8, these most connected genes are considered as hub genes, including *map3k10*, *cblb*, *nck2*, *nin*, *plk2b* (LOC103398298), *prkag1*, of which *plk2b* was also the common genes in five comparisons (Table 3).

**Fig. 8.**
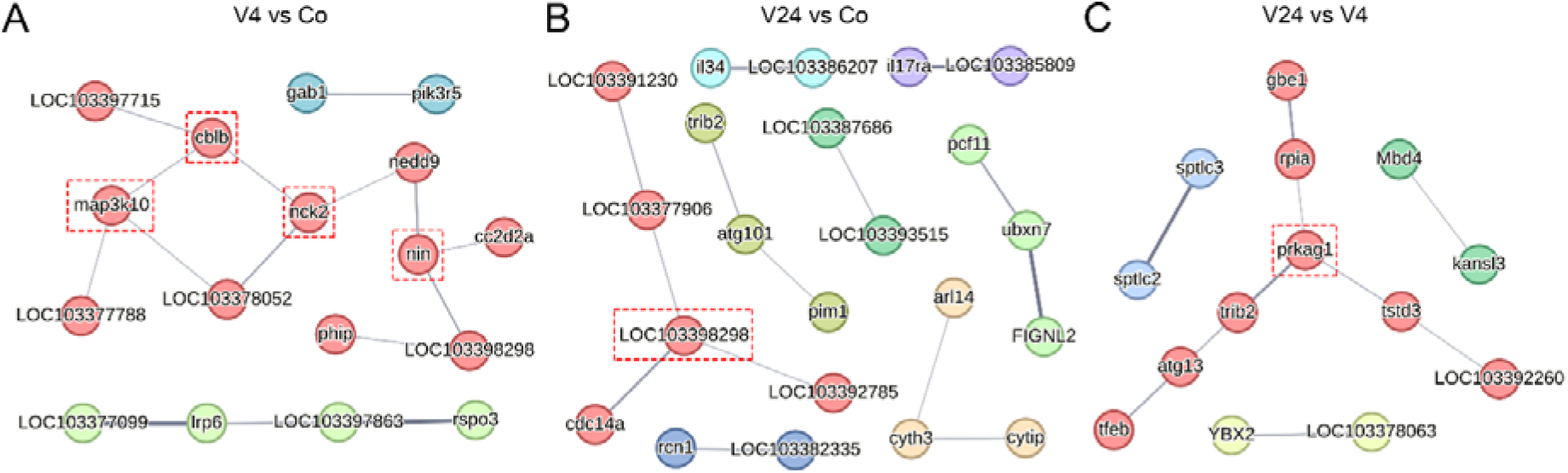
Protein-protein interaction (PPI) showing the hub common genes in the comparison of V4 vs Co (A), V24 vs Co (B), and V24 vs V4 (C). Co, control; V4, 4 h after the infection; V24, 24 h after the infection.

### 3.6. Representative gene analysis

One immune-related gene, *peli2*, that underwent both significant differential m6A methylation and differential expression was selected for the visualization of read mapping depth and for the validation of significance intuitively. The gene *peli2* was hyper m6A methylated and up-regulated in the comparison of V4 vs Co. As shown in Figure 9, the levels of gene expression (input) and the m6A methylation (IP-input) are markedly higher at V4 group compared with the control, considering that the overall sequence depth are similar between groups.

**Fig. 9.**
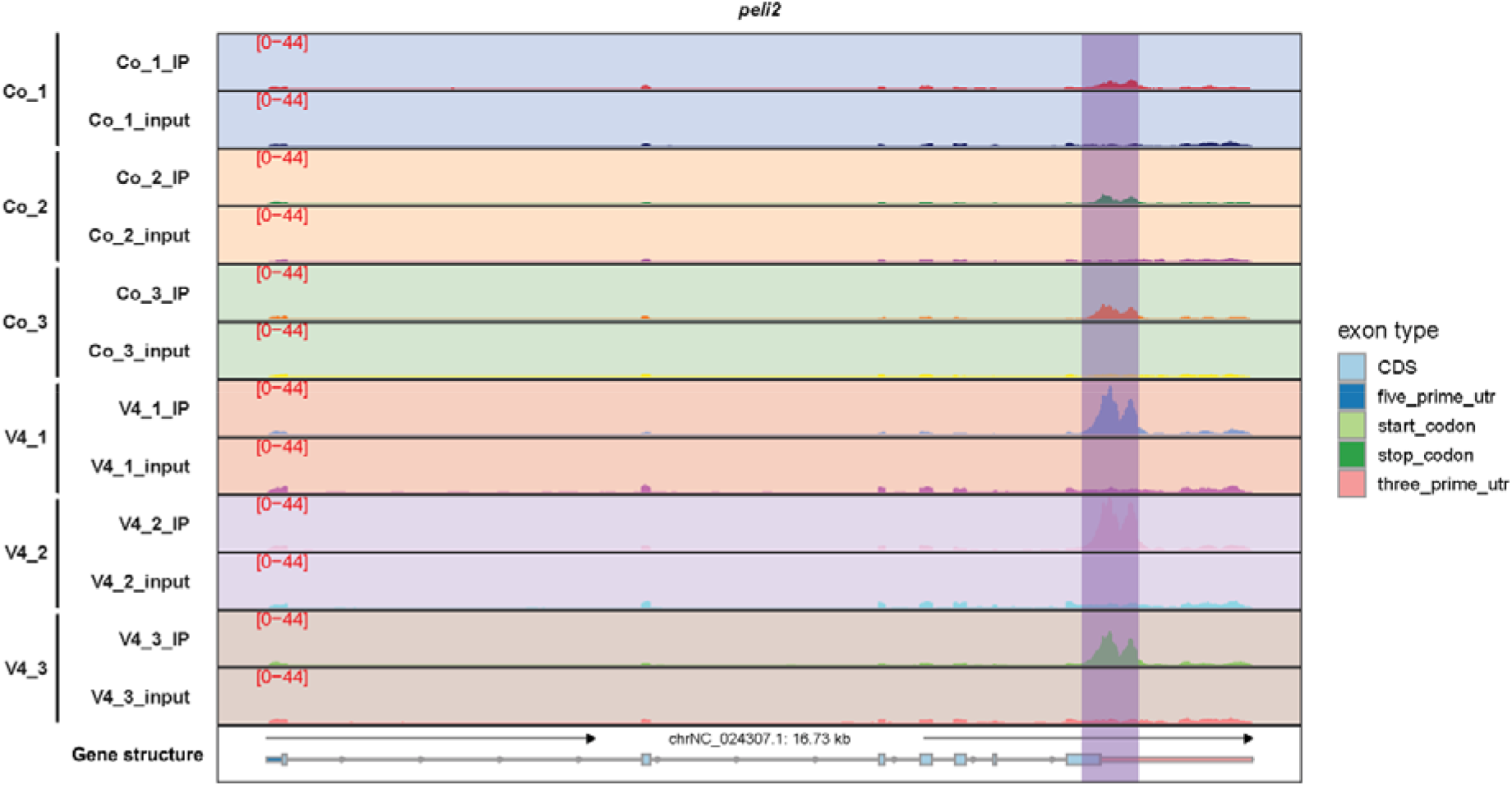
Visualization of the read mapping depth for one common gene of DMG and DEG, *peli2*, in the comparison of V4 vs Co. Co, control; V4, 4 h after the infection.

### 3.7. qRT-PCR validation

To validate the accuracy of RNA-seq analysis, qRT-PCR was performed. As shown in Figure 10, two sets of data show good consistency, i.e., the trend (up-regulation or down-regulation) and the magnitude of change are comparable between RNA-seq and qRT-PCR. qRT-PCR successfully validated the RNA-seq analysis, strengthening the confidence in the data obtained from the study.

**Fig. 10.**
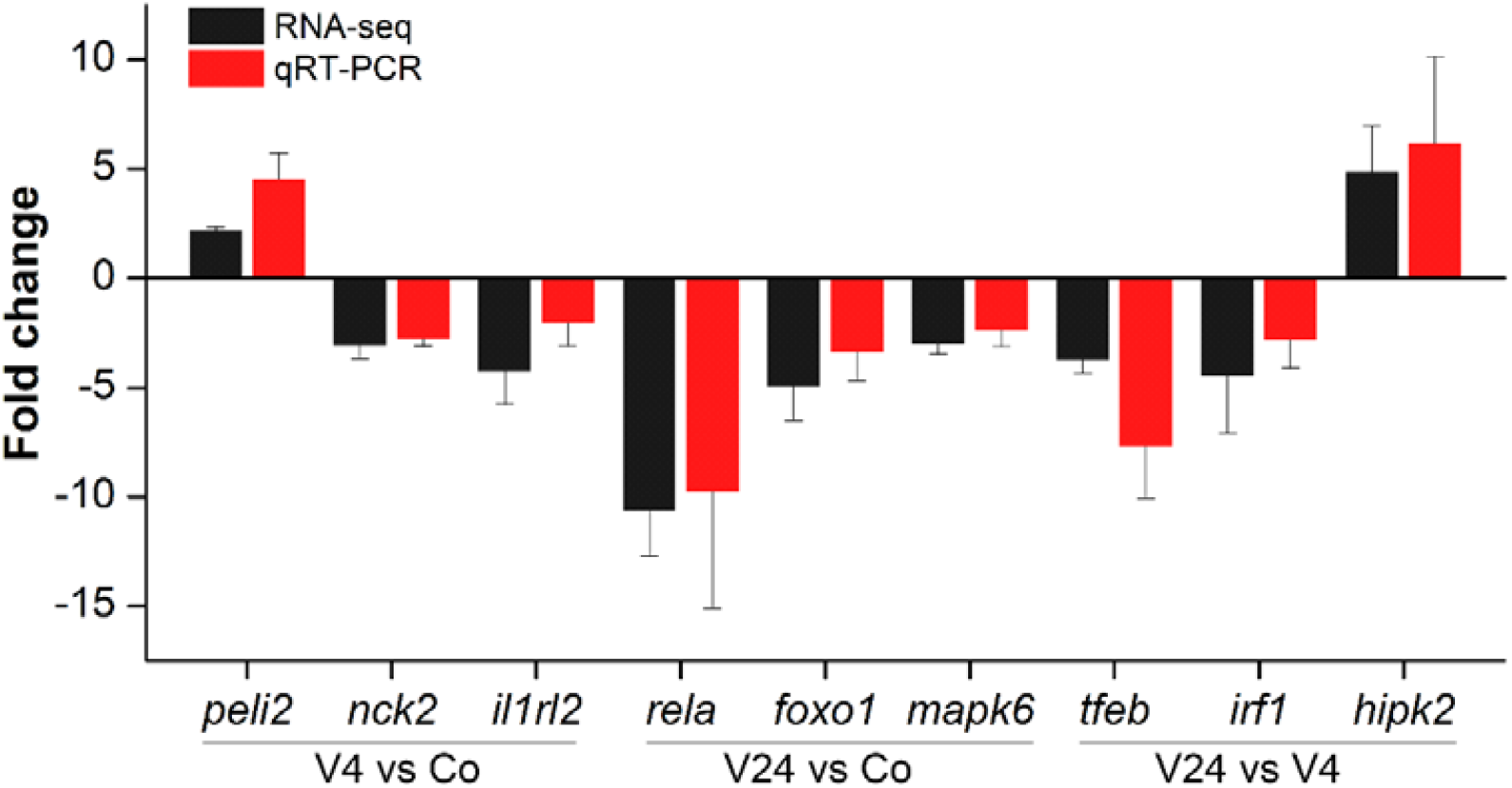
Validation of the expression of common differential genes using quantitative real-time PCR. Fold change value indicates the relative expression. Co, control; V4, 4 h after the infection; V24, 24 h after the infection. *peli2*, pellino E3 ubiquitin protein ligase family member 2; *nck2*, NCK adaptor protein 2; *il1rl2*, interleukin-1 receptor-like 2; *rela*, RELA proto-oncogene, NF-kB subunit; *foxo1*, forkhead box O1; *mapk6*, mitogen-activated protein kinase 6; *tfeb*, transcription factor EB; *irf1*, interferon regulatory factor 1; *hipk2*, homeodomain interacting protein kinase 2.

## 4. Discussion

The m6A modification plays a pivotal role in regulating gene expression in many aspects, and subsequently, diverse biological processes including immune response (He and He, 2021). These knowledge have been mainly gained through mammalian research, but whether or not and to what extent m6A is involved in aquaculture fish species is underexplored. In this study, we present an intriguing exploration of the characterization of m6A methylation and its involvement in modulating the immune response of an important aquaculture species, *C. semilaevis*, against the pathogenic bacterium *V. anguillarum*. For the first time, we characterized the number, distribution, and motif of m6A in genes of *C. semilaevis*, as well as revealed its dynamic implication and potential functions in gene expression during the continuous response following the bacterial infection. A set of candidate genes and enriched pathways, not only the immune-related ones, were identified worth further analysis. This study not only adds to our understanding of the complex regulatory mechanisms involved in fish immune responses, but also opens new avenues for disease control in aquaculture.

The identification of over 20,000 m6A peaks in over 12,000 genes suggests a widespread presence of m6A modifications in the transcriptome of *C. semilaevis*, aligning with the findings in other species which conclude that the m6A modification is a prevalent post-transcriptional modification in eukaryote (Dominissini et al., 2013; Meyer and Jaffrey, 2014). The analysis of peak distribution on RNA functional elements reveals that CDS contains the highest number of peaks, followed by stop codon, 3’ UTR, and start codon. The observation of markedly elevated peak density in the vicinity of start codon and stop codon, with the latter more pronounced than the former, is consistent with studies in other species (Dominissini et al., 2013; Dominissini et al., 2012; Meyer and Jaffrey, 2014; Meyer et al., 2012), indicating specific functions of m6A in these regions, such as affecting translation initiation and termination. In fact, recent studies have revealed that the m6A “writer” METTL3 increased the translation of mRNAs when it was tethered to sites near stop codon (Choe et al., 2018).

Our motif analysis provided insights into the sequence characteristics associated with m6A modification by the methyltransferase across different time points following infection (Fig. 2). As expected, the canonical DRACH motif was enriched in samples at all three time points (0, 4, and 24 hpi) (Fig. 2A, F, G). This finding reinforces the established role of the DRACH motif as a primary recognition element for m6A methyltransferase binding (Linder et al., 2015; Wang et al., 2021). Not only that, our analysis extends beyond the canonical motif by identifying additional enriched motifs specific to each time point. Interestingly, motifs containing GAA emerged as significantly enriched compared to the background at all three time points (Fig. 2B, 2E, and 2I), suggesting a potential role for the GAA sequence in m6A modification. The enrichment of the different motif observed at different time points after infection suggests a potentially unique regulatory landscape for m6A deposition during the early stages of infection. Further investigation into the functional roles of these motifs and their potential interplay with trans-acting factors will provide a more comprehensive understanding of the dynamic regulation of m6A during infection and other biological processes.

One of the key findings of this study is the identification of a set of differentially m6A methylated genes that are associated with the immune response against *V. anguillarum*, including three common DMGs, including *pdp1*, *rgs5l*, and one uncharacterized gene (LOC103388921), across all three comparisons. These genes may represent key regulators of the host response to bacterial infection, whose functions are modulated by m6A modifications. The *pdp1* gene encodes pyruvate dehydrogenase phosphatase catalytic subunit 1, which activates the pyruvate dehydrogenase complex to convert pyruvate into acetyl-CoA for ATP and energy production (Gray et al., 2014). It is not surprising that energy-related genes were regulated. Energy production and allocation are crucial for effective immune response (Wang, A. et al., 2019). Various immune-related processes, including immune cells generation/differentiation, antibodies production, and inflammatory responses mounting, are energy-intensive. In addition, in the face of immediate threats such as infection, the body may divert energy from other functions to support the immune response. Noting that in addition to *pdp1*, the 11 most common differential genes among the six comparisons of DMGs and DEGs (Table 3) include another two energy-related genes, i.e., *cyth3* and *slc16a2*, highlighting the importance of energy metabolism in immune response. The *rgs5l* gene encodes regulator of G-protein signaling 5-like. Rgs5 has been shown to facilitate the production of cytokines and inflammation via tumor necrosis factor (TNF) signaling (Yin et al., 2023). Furthermore, it determined neutrophil migration in the acute inflammatory phase of lung injury (Sharma et al., 2021). This immune-related gene under m6A modification may be important for the immune response in this study. Future studies exploring the biological roles and mechanisms of these genes may provide valuable insights into host immunity.

In addition to the study of m6A methylation, we also performed analysis on gene expression. The series test of cluster analysis performed in this study has provided a comprehensive overview of the dynamic changes in gene expression following bacterial infection. The clustering of genes into eight groups with distinct expression patterns highlights the complexity of the host response to such an infection. For instance, we observed an initial down-regulation and the subsequent up-regulation in gene expression of clusters 1, 5, and 8, albeit to varying degrees. There are likely two reasons for this phenomenon. The host may effectively respond to the infection by down-regulating the expression of genes involved in certain pathways to reallocate resources and energy to immune responses (Wang, A. et al., 2019). In addition, the down-regulation of certain genes may be not the result of active host regulation, but caused by the negative effects of the bacterium where they release toxins to help them survive and proliferate within the host cells (Bhavsar et al., 2007). Moreover, the overall up-regulation of gene expression in cluster 7 suggests a sustained activation of specific immune pathways that are important for the host to control the disease. Genes with expression trend, especially those with significant changes, could serve as potential biomarkers for disease diagnosis/progression and as novel therapeutic targets.

Our analysis revealed a striking enrichment of the FoxO signaling pathway across all six experimental comparisons (Table S3). This finding suggests a central and ubiquitous role for FoxO signaling in the immune response examined in this study. In addition to their key roles in cellular survival, apoptosis, and metabolism, FoxO transcription factors have been shown to play critical roles in stress resistance and the homeostasis of immune-relevant cells, including T cells, B cells, and non-lymphoid lineages (Peng, 2008). In human, *Drosophila,* and shrimp (*Marsupenaeus japonicus*), FoxO signaling could activate the production of antimicrobial peptides (AMPs) to fight pathogen infections, where the regulation was evolutionarily conserved (Becker et al., 2010; Fink et al., 2016; Li et al., 2021). Furthermore, FOXO members have been found to be important in aquaculture fish species after bacterial infections, including largemouth bass (*Micropterus salmoides*) (Zhou et al., 2021), turbot (*Scophthalmus maximus*) (Pan et al., 2021), and channel catfish (*Ictalurus punctatus*) (Gao et al., 2019). For instance, FoxO3 was found to modulate LPS-activated hepatic inflammation in turbot. Together with these valuable previous analyses, this study suggests that FoxO activation is a key component of the cellular response to bacterial infection in fish species. Functional studies, such as siRNA knockdown or overexpression of key members in FoxO family and the signaling pathway, could elucidate the specific contribution of FoxO signaling to the immune response. In addition, six pathways were significantly enriched in five comparisons (Table S3), including MAPK signaling pathway, neuroactive ligand-receptor interaction, base excision repair, PPAR signaling pathway, mRNA surveillance pathway, endocytosis, suggesting their importance in the immune response.

Besides the FoxO signaling pathway, the base excision repair (BER) pathway is the only pathway that is enriched in all three DMG comparisons, and it is also the commonly enriched pathway in two DEG comparisons (V24 vs V4, V4 vs Co). The BER pathway is primarily responsible for removing and repairing base lesions caused by oxidation, depurination, alkylation, and deamination, which help maintain the integrity of the genome under constant threat of environmental stresses (Stratigopoulou et al., 2020). Keeping genome integrity in immune cells is extremely important for their homeostasis, differentiation, and proliferation in response to internal and external stresses such as infection (Zhao et al., 2022). Except for its functions in DNA repair, BER pathway has been recently shown to regulate RNA metabolism and gene expression through transcriptional and post-transcriptional mechanisms (Antoniali et al., 2017), indicating this pathway may be a regulatory all-rounder.

Differential m6A methylation and expression analysis showed that *C. semilaevis* could exert timely transcriptome-wide regulation responding to the *V. anguillarum* infection as early as 4 h post infection. This is a novel finding, and further exploration is needed to determine whether these regulations could occur earlier after infection. Importantly, our joint analysis of DMGs and DEGs provides valuable insights into the dynamic regulation of gene expression during bacterial infection. The Venn diagrams (Fig. 7A, E) reveal a substantial proportion of DMGs that could regulate differential gene expression at both the early stage (V4 vs Co) and the later stage (V24 vs V4), suggesting potent regulatory roles of m6A methylation on gene expression regulation. The effect of m6A on gene expression in this study appears to be context-dependent, with evidence for both down-regulation and up-regulation depending on the stage of bacterial infection. m6A modification was shown to mainly reduce gene expression during the early stage (0-4 hpi) of bacterial infection. However, during the later stage (4-24 hpi), m6A modification is associated with increased expression of specific genes. This observation suggests a potential shift ability of m6A in dynamically regulating gene expression. Future studies should explore the functional roles of the common genes in all comparisons and decipher the mechanism underlying gene expression regulation mediated by specific m6A methylation.

Furthermore, the combined analysis of DMGs/DEGs and PPI analysis reveal a single candidate gene, *plk2b*, indicating its importance in immune response of this study. The family of Polo-like kinases (PLKs) are evolutionary conserved from yeast to human, and PLK2 is extensively involved in normal physiological functions, stress responses to external stimuli, and diseases progression (Zhang et al., 2022). In LPS-induced inflammation, expression of *plk2* is elevated, which phosphorates a disintegrin and metalloprotease 17 (ADAM17) and results in the release of pro-TNFα from primary macrophages and dendritic cells. In addition, PLK2 is essential for viral sensing *in vitro* and *in vivo* in immune dendritic cells of mice (Chevrier et al., 2011). Zhou et al. (Zhou et al., 2021) found that *plk2* may play an important role in the immune system response of largemouth bass infected by *A. hydrophila*. In this study, the expression of *plk2b* gene was first significantly down-regulated (V4 vs Co) accompanied by hyper-m6A methylated and then up-regulated (V24 vs V4), and overall, the gene was cataloged into “hyper-down” (V24 vs Co), suggesting the importance of this gene and the dynamic role of m6A on gene expression.

It is worth noting that while this study has made significant progress in understanding the role of m6A methylation in fish immune responses, there are still many unanswered questions. For example, it remains unclear how m6A methylation targets and regulates immune-related genes, whether the immune response could be enhanced by manipulating the expression levels of important enzymes (“writers,” “erasers,” and “readers” of m6A methylation) and identified candidate genes. Future studies should aim to address these questions by using more tissues/time points and employing more advanced techniques to gain a deeper understanding of the spatiotemporal molecular mechanisms underlying m6A-mediated immunoregulation.

## 5. Conclusions

This study provides valuable insights into the epitranscriptomic regulation of immunity in fish, which is an underexplored area compared to mammals. For the first time, we reveal the rapid and dynamic modulation of m6A methylation in the immune response against *V. anguillarum* in half-smooth tongue sole. Key genes and pathways were identified, including immune-related and energy metabolism-related ones, emphasizing their importance in combating bacteria. This study not only adds to our knowledge of the complex regulatory networks involved in immunity but also opens up new possibilities for disease control in aquaculture. Further functional analyses are warranted to formulate novel and optimal strategies for the sustainable development of aquaculture.

## CRediT authorship contribution statement

**Suxu Tan**: Conceptualization, Funding acquisition, Experimental operation, Data analysis, Writing - original draft, review and editing. **Wenwen Wang**: Experimental operation, Manuscript revision. **Sen Han**: Experimental operation. **Kunpeng Shi**: Manuscript revision. **Shaoqing Zang**: Manuscript revision**. Zhendong Wu**: Experimental operation. **Zhenxia Sha**: Supervision, Funding acquisition, Manuscript revision.

## Declaration of Competing Interest

No potential conflict of interest was reported by the authors.

## Acknowledgment

This study was supported by the National Key R&D Program of China (Grant No. 2022YFD2400401), Natural Science Foundation of Shandong Province (Grant No. ZR2023QC259), Shandong Key R&D Program (For Academician team in Shandong (Grant No. 2023ZLYS02), and Taishan Scholar Youth Project of Shandong Province, China.

## Notes

### Competing Interest Statement

The authors have declared no competing interest.

